# Application of Volumetric Absorptive Micro Sampling to Measure Multidimensional Anti-Influenza Hemagglutinin Igg Antibodies by MPlex-Flu Assay

**DOI:** 10.1101/588038

**Authors:** Jiong Wang, Dongme Li, Alexander Wiltse, Jason Emo, Shannon P. Hilchey, Martin S. Zand

## Abstract

Recently, volumetric absorptive microsampling (VAMS) has been used for peripheral blood sampling and analyses in several fields. VAMS ensures accurate sampling by collecting a fixed blood volume (10 or 20 *µ*L) on a volumetric swab in blood spot format, and allows for long-term sample storage. The mPlex-Flu assay is a novel, multidimensional assay that measures the concentration of antibodies against multiple influenza virus hemagglutinins simultaneously strains with a small volume of serum (less than 5 *µ*L). Here we describe combining these two methods to measure multidimensional influenza antibody activity using a finger-stick and VAMS. In this study, we compared influenza antibody profiles measured from capillary blood obtained with a finger-stick, and venous whole blood collected by traditional phlebotomy from 20 subjects using the mPlex-Flu assay. We found that results with the two sampling methods were virtually identical across all influenza strains within the same subject (mean of *R*^2^ =0.9470), and that antibodies remained stable over three weeks when VAMS samples were stored at room temperature and transported using a variety of shipping methods. Additionally, VAMS sampling is an easy and highly reproducible process; when volunteers performed finger stick VAMS at home by themselves, the results of anti-HA antibody concentrations showed that they are highly consistent with sampling performed by study personnel on-site (*R*^2^ =0.9496). This novel approach provides advantages for clinical influenza vaccine studies, including ease of sampling, low cost, and high accuracy. We conclude that these methods could provide an accurate and low-cost means for monitoring the influenza virus antibody responses in large population studies.

## 1 Introduction

Both seasonal and emerging influenza virus infection are among the largest reoccurring global public health threats [1], and vaccination is the major method of prevention[2]. Flu vaccines are currently designed to elicit antibodies against hemagglutinin (HA), the most abundant glycoprotein on the viral surface[3]. Protective antibodies block the ability of HA to bind to sialic acid on target cells, or enhance viral clearance, preventing infection [4]. Measuring antibody-mediated immunity is critical to evaluate the preventing immunity to seasonal and emerging influenza viruses. However, obtaining serum samples to measure antibody-mediated influenza immunity is a resource intense, time consuming, and expensive process[5]. This limits our ability to conduct large-scale influenza vaccine clinical trials, measure population immunity, and assess the mismatch between circulating influenza strains and the seasonal influenza vaccine in real time. Several factors contribute to this issue. Most assays of antibody-mediated influenza immunity, such as hemagluttinin inhibition (HAI), enzyme linked immunosorbent assay (ELISA), and microneutralization (MN) assays all require at the minimum 0.1 - 0.2 mL of serum to perform with appropriate replicates. Obtaining such quantities of serum is done by venupuncture phlebotomy performed by a healthcare professional, requiring subject travel to the research facility or collection point. Finally, blood samples require post-phlebotomy processing including serum separation, aliquotting, and storage. Thus, developing a solution would require both a simple method for in-the-field collection of small amounts of peripheral blood or serum, coupled with an assay that uses very small sample sizes (10-20 *µ*L). Here we describe and validate such a system, using a combination of volumetric absorptive microsampling (VAMS)[6] coupled with a Luminex-based assay (mPLEX-Flu)[7, 8, 9, 10, 11, 12] to quantitatively measure IgG antibodies against 33 strains of influenza hemagglutinin.

We have previously developed a Luminex-based multiplex assay (mPlex-Flu assay) that can simultaneously measure absolute antibody concentrations (IgG, IgM or IgA) against up to 50 influenza strains using ≤ 5*µL* of serum[7, 8, 11, 12]. The mPLEX-Flu assay has a continuous linear read-out over 4 logs, with low Type-I (false positives, specificity) and Type-II (false negatives, sensitivity) errors[10]. It provides absolute concentrations for strain-specific anti-influenza IgG antibody levels, as opposed to 8-12 discrete titer levels for other assays (e.g. HAI, MN), with extremely low inter- and intra-subject variance [9]. Notably, the mPlex-Flu assay also has a very high correlation with both standard HAI and MN assays, with several added advantages, including simultaneous measurement of absolute anti-HA IgG levels for a large number of influenza strains[7, 8, 11, 12], greater precision of clinical trial group statistical comparisons[9, 10], and a low per-sample cost.

Development of a method to measure anti-HA IgG levels using small volume blood samples simply and remotely collected by study subjects would greatly improve our ability to conduct more robust clinical trials, population immunity surveys, and augment current influenza surveillance efforts. For example, most influenza vaccine clinical trials have measured anti-HA IgG titers in peripheral blood samples pre-vaccination (day 0), and at days 7 and 28 post-vaccination[13], while others have also collected samples at time more distant time points[14]. A substantial expense in these trials is sample collection by trial personnel. For the same reasons (i.e. cost and inconvenience of phlebotomy for sample collection), large-scale surveys of population antibody-mediated immunity to influenza are rare. We are not aware of any current studies assessing IgG mediated immunity to multiple influenza strains with over 1,000 subjects. Finally, the the United States Center for Disease Control and the World Health Organization both conduct extensive influenza virus field surveillance programs[15], collecting viral samples by nasal swab to isolate and sequence influenza strains in people with influenza-like illnesses. Yet, these programs generally do not collect serum to assess antibody-mediated influenza immunity, likely due to the cost and time needed for proper phlebotomy and sample processing. In all three of these examples, a simple method to collect and analyze serum samples for anti-influenza IgG would decrease the barriers to multiple sample collection (cost, inconvenience, sample processing), and improve scientific knowledge of influenza immunity.

One approach to simplifying sample collection is to perform a finger or heal stick to draw a drop of blood, using a disposable lancet as is done for diabetic blood glucose monitoring. The blood drop, generally 50-200*µ*L, is then adsorbed onto filter paper and dried. Samples are then eluted and analyzed at a later date. This micro-sampling to dried blood spot (DBS) method was first introduced in 1963[16]. It has been used to assess the HIV-1 antibodies in newborns, in population-based surveys for more than 25 years[17, 18, 19], and for analysis of anti-drug antibodies in FDA clinical trials. DBS is safer and simpler than venupuncture. It enables self-sampling at home and can greatly reduce costs for clinical or population-based studies. A significant drawback of DBS, however, is the high variability of sample volumes. This makes calculation of a concentration problematic and limiting its use for quantitative measures of antibody abundance. In contrast, volumetric absorptive microsampling (VAMS) devices adsorb a consistent volume of blood from a finger-stick, generally 10 or 20*µL*, and have been recently used to collect samples for antibody testing in many fields (Reviewed in[6]). This new technique overcomes the issue of inconsistent blood volumes between sample blood spots in the DBS method. VAMS allows accurate and precise sampling with standard deviation ≤ 0.4*µL* with 10*µL* blood samples[20]. In addition, IgG and IgM antibodies in the dried blood sample are known to be stable at room temperature for weeks, and at −20°C for years.

Here we describe using a combination of VAMS with mPlex-Flu to quantitatively measure IgG antibodies against 33 strains of influenza virus HAs. This study validates the the accuracy, reproducibility, and sample stability of this novel assay combination. Overall, it provides direct evidence that application of VAMS in mPlex-Flu assay will be a powerful tool to generate extensive samples and high dimensional data concerning influenza strain-specific antibody mediated immunity for use in influenza vaccine and population immunity studies.

## 2 Material and Methods

### 2.1 Human Subjects Ethics Statement

This study was approved by the Research Subjects Review Board at the University of Rochester Medical Center (RSRB approval number RSRB00070463), and informed consent was obtained from all participants. Research data were coded such that subjects could not be identified, either directly or through linked identifiers. Subject identification numbers were re-encoded for publication.

### 2.2 Participants and Sample Collection

Twenty one healthy volunteers 18-65 years of age were recruited for this study. Subjects who were taking immunosuppressive medications were excluded. All subjects had samples collected by both venous phlebotomy and VAMS (Figure 1).

**Figure 1.**
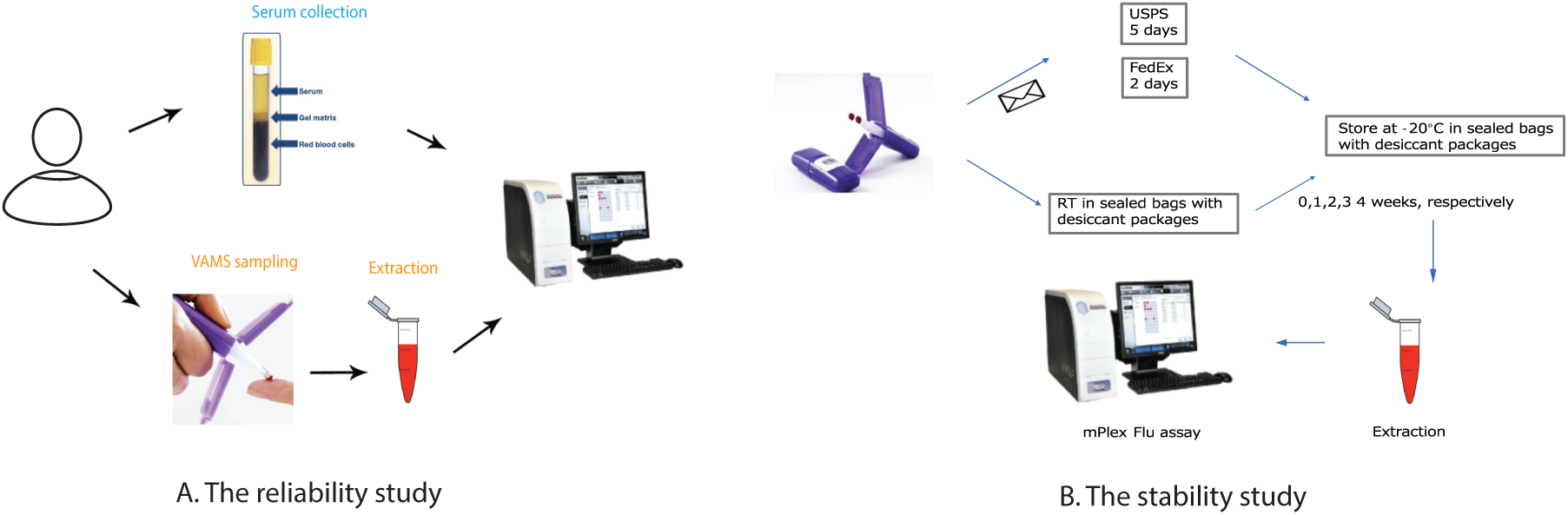
The experimental design. The experiments were conducted in 2 sets (A) A reliability study where parallel samples from phlebotomy (serum) and VAMS sampling serum, which were then analyzed by mPLEX-Flu and compared. (B) The stability study involved multiple samples from the same donor that were then either left at room temperature for

For venous phlebotomy, standard venupuncture was performed and 3-4 mL of blood was collected in a serum collection tube (BD, NJ, USA). Samples were immediately centrifuged (3000 RPM, 4°C, 12 minutes), aliquotted into 100*µL* cryo-vials, and stored at −20°C until analysis.

### 2.3 Study strategy

The study was designed to assess both variability between standard venupuncture for serum and VAMS sampling, and reproducibility of results when subjects performed VAMS sampling remotely after instruction. At the initial study visit, each volunteer donated one venous blood sample by phlebotomy, and one VAMS blood sample by finger-stick. Both samples were collected by study coordinators on site at the University of Rochester Clinical Research Center. Study subjects were then trained to perform a finger-stick with the lancet device, and collect the VAMS sample. And after training, one VAMS kit was sent home with the volunteer. Three days later, the volunteer self-collected a second VAMS sample and returned it in sealed packaging to the Research Center for analysis.

VAMS blood samples were collected using the manufacturer’s 10*µL* collection kit (Mitra Collection Kit; Neoteryx, CA, USA). After alcohol swabbing, the lateral portion of the participant’s finger was punctured using a contact-activated lancet. Gentle pressure was applied to the finger to allow a drop of blood to collect at the skin surface. A porous, hydrophilic VAMS tip was held against the blood drop until completely filled with blood, which took around 2-3 seconds. Each tip absorbed 10*µL* of blood. There were two tips in each collection kit, for a total of 20*µL* of blood per collection. Tips were allowed to dry for 1-2 hours at room temperature in protective cassettes that prevented them touching each other or their surroundings.

For the stability study, one donor’s 14 VAMS blood samples were collected at the same time using the same method.

### 2.4 Storage of VAMS samples

All VAMS tips were placed in separate and sealed containers with silica desiccant packets, and all venous phlebotomy serum samples were stored in sealed cryo-vials. Both types of samples were stored at −20°C until analysis.

### 2.5 Extraction of antibodies from VAMS samples

The absorbent tips from each VAMS collection kit (containing 10*µL* blood sample) were soaked in 200*µL* extraction buffer (PBS + 1% BSA + 0.5% Tween) in 1 mL 96 well plates (Masterblock, GBO, Austria) and shaken overnight to extract the antibodies as described previously[19].

### 2.6 mPlex-Flu assay

The mPlex-Flu assay was performed as described previously[7, 8, 12]. Briefly, venous phlebotomy serum samples were diluted 1:5000 with PBS, while the 200*µL* extractions from VAMS device (1:200) were further diluted 1:250, to yield a final 1:5000 dilution of the VAMS samples. For both serum and VAMS samples, 200*µL* of diluted sample was used for analysis and added to a black, clear-bottom 96 well plate (Microplate, GBO, Austria). Standard serum (STD01) was made in our laboratory[7, 12], and the standard curve for each influenza virus strain was generated by 1:4 serially diluting STD01 for each batch of samples. 50*µL* of the diluted sample was added into each reaction well. All samples were run in duplicate.

The influenza HA bead panel used in this study is shown in Table 1, comprising 33 separate rHAs. 50*µL* of bead mix was added to each well of the plate as previously described[7, 8, 12]. Plates were then incubated with gentle shaking for 2 hours at room temperature, and then washed (PBS + 1% Bris + 0.1% BSA). A magnet placed under the plate immobilized the beads during washes. After three washes, a goat anti-human IgG-PE secondary antibody (Southern Biotech, Cal No:2040-09) was added, and plates were incubated for another 2 hours. After three more washes, beads were re-suspended in drive fluid (Luminex Co., TX) and the beads were analyzed using MAGPIX^™^ Multiplex Reader (Luminex Co., TX). The calculation of IgG antibody concentration against each individual influenza virus strain rHA was performed by Bio-Plex Manager^™^ 6.2 software (Bio-Rad Co., CA).

**Table 1.**
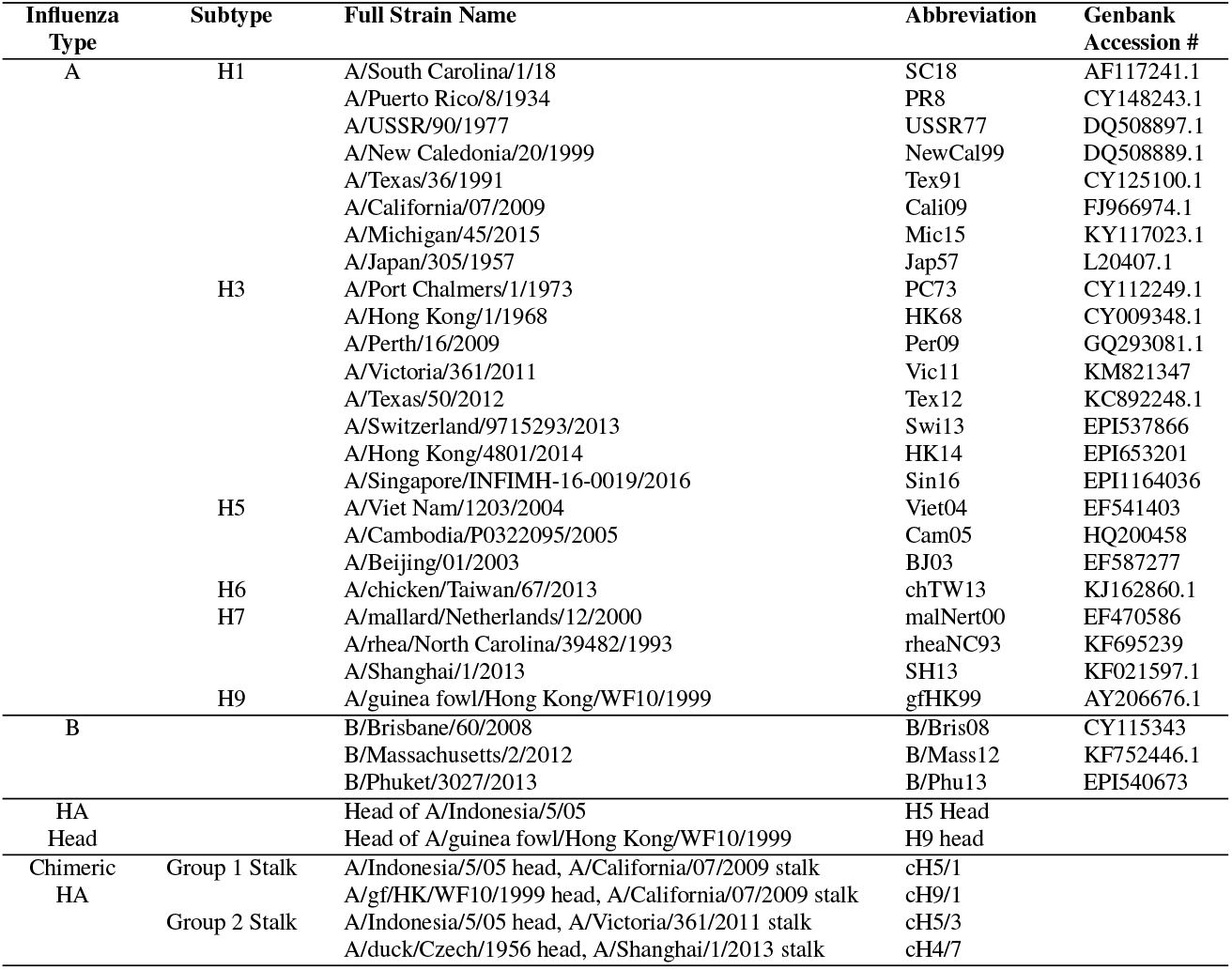
The panel of influenza virus rHAs used in mPlex-Flu assay

### 2.7 VAMS sample stability analysis

To assess the stability of VAMS samples at room temperature over time, 14 VAMS samples were collected from the same subject. An initial 2 VAMS samples were stored at −20°C after drying for 2 hours. A further 8 VAMS samples were left on the lab bench at room temperature (22-25°C, controlled, but not monitored). Two of these were then moved into 20°C storage at days 7, 14, 21 and 28 after the initial sampling. The remaining 4 VAMS samples were mailed back to the lab using the United States Postal Service and 2-day overnight delivery. The samples sent by 2-day overnight delivery returned in 2 days and those sent by the United States Postal Service returned in 5 days (Figure 1). The VAMS samples were stored at −20°C upon arrival in the laboratory.

### 2.8. Statistical analysis

Spearman’s correlation coefficient[21] was used to measure the reliability of mPLEX-Flu results from VAMS versus conventional venous phlebotomy samples, the reproduciblity of mPLEX-Flu results from VAMS collected by volunteers at home verses VAMS collected by study coordinators on-site, and the stability mPLEX-Flu results from VAMS samples stored at room temperature over time or after shipping. For calculation of correlation coefficients, measurements from the mPlex-Flu assay using various VAMS samples were either combined across multiple influenza virus types or separated by influenza virus type and subject.

Because the sample size is small and the data were not normally distributed, we used generalized linear models with an identity link function[22] to compare the mean measurements from the mPlex-Flu assay results obtained with VAMS versus conventional serum sampling under different room temperature storage times and shipping methods. Statistical analysis software SAS v9.4 (SAS Institute, Inc., Cary, NC) and R version 3.5.1 were used for all the data analysis. The significance level for all tests was set at p=0.05.

## 3 Results and discussion

### 3.1 Subject demographics

Twenty one healthy volunteers were recruited for this study and their demographics are shown in Table 2. More female subjects took part in this study (67%) than male, with majority of volunteers being Caucasian. The distribution of age groups is relatively uniform with fewer volunteers age 20 or less.

**Table 2.** Subject Demographics

|  | Summary | N (%) |
| --- | --- | --- |
| Sex | Female | 15 (67) |
|  | Male | 6 (33) |
| Ethnicity | Hispanic / Latino | 0 (0) |
|  | White | 19 (90) |
|  | Black or African American | 0 (0) |
|  | Asian | 2 (10) |
|  | Native American or Alaska Native | 0 (0) |
|  | Native Hawaiian or other Pacific Islander | 0 (0) |
| Age | <20 | 2 (10) |
|  | 21-45 | 7 (33) |
|  | 46-60 | 7 (33) |
|  | 61-75 | 5 (24) |
| Total |  | 21(100) |

### 3.2 mPlex-Flu assay results from VAMS and serum sampling are highly correlated

In order to compare the variability of mPLEX-Flu testing of capillary blood VAMS versus venous serum sampling, we evaluated 20 participants who each coincident VAMS finger stick and venupuncture blood collection, with mPlex-Flu assay on both sampling types. The resulting data are shown in heatmap form in Figure 2. We then calculated the ratio of mPLEX-Flu values from VAMS sampling as a fraction of that from serum for each strain:

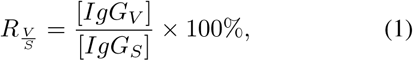

where *V* denotes a VAMS sample and *S* denotes a serum sample.

**Figure 2.**
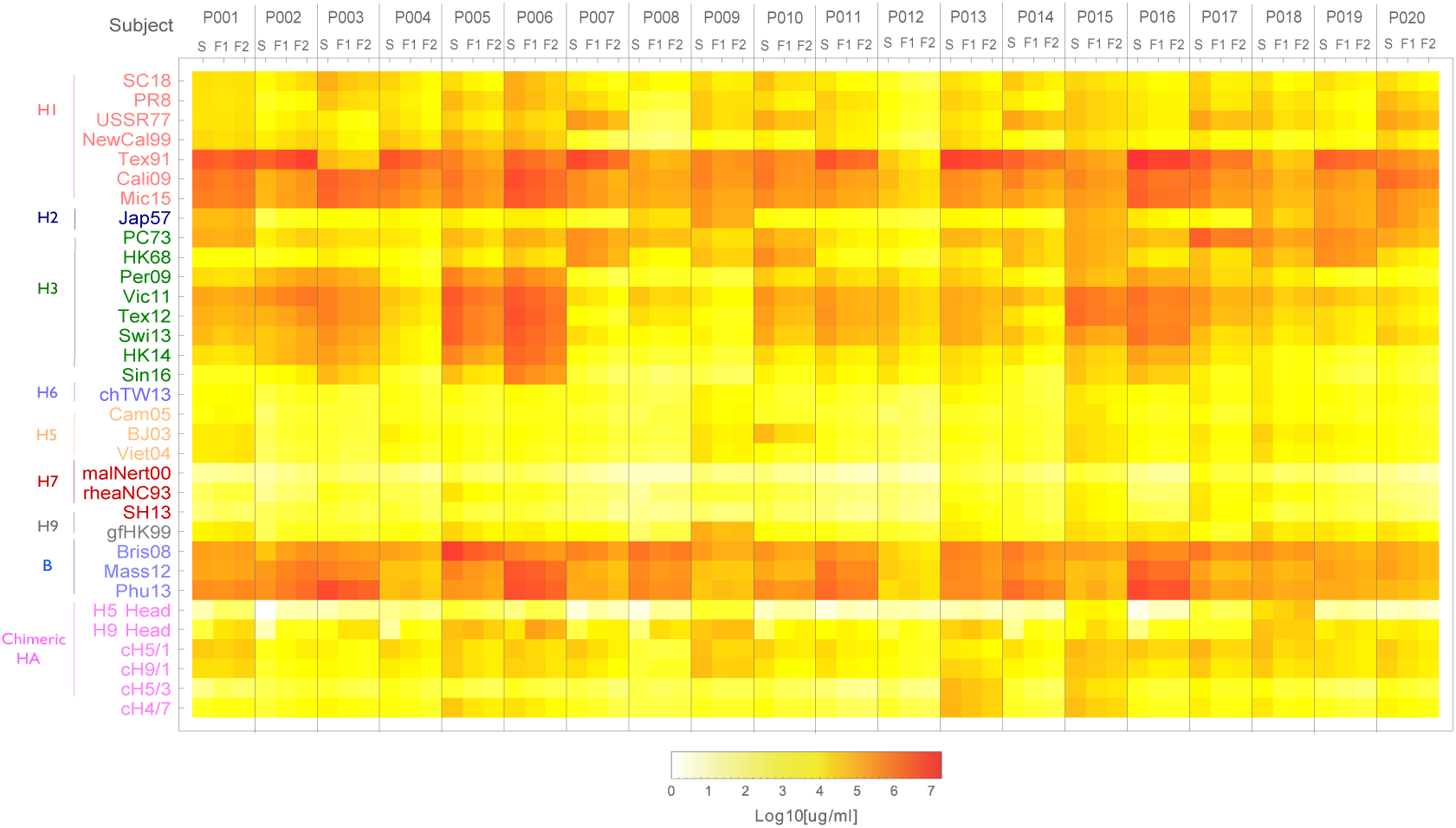
The IgG antibody concentration against 33 individual influenza virus strains assessed by mPlex-Flu assay in this study. The blood samples of twenty subjects were collected trough phlebotomy serum sampling(S), VAMS sampling on-site (F1), and VAMS sampling at-home (F2) were testing by mPlex-Flu assay with a 33 influenza virus HAs panel in the same dilution. The IgG concentrations of samples were calculated based on a standard curve for individual virus strain generated by standard serum with Bio-Plex Manager^™^ 6.2 software. The Mean concentration of duplicate were shown in the heat-map (details see the material and methods section).

To compare the anti-HA IgG concentrations from mPlex-Flu in samples obtained by VAMS versus traditional phlebotomy, we used the Spearman’s Correlation Coefficient[21] with the Benjamini-Hockberg multiple testing correction method[23]. All data are listed in Table 3. There was only modest variation (*Ratio*) of the anti-HA IgG antibody concentrations in mPlex-Flu assay of VAMS sampling (*V AMS*) versus conventional serum sampling (*Ration* from 108 to 112%). Comparing the ratio *Ration_*V AMS/SER*_* for each influenza virus strain, the anti-HA antibody concentrations from VAMS are highly correlated with those from serum sampling (P<0.0001). These results suggested that VAMS sampling can generate almost identical and highly correlated antibody concentration data in the mPlex-Flu assay.

**Table 3.**
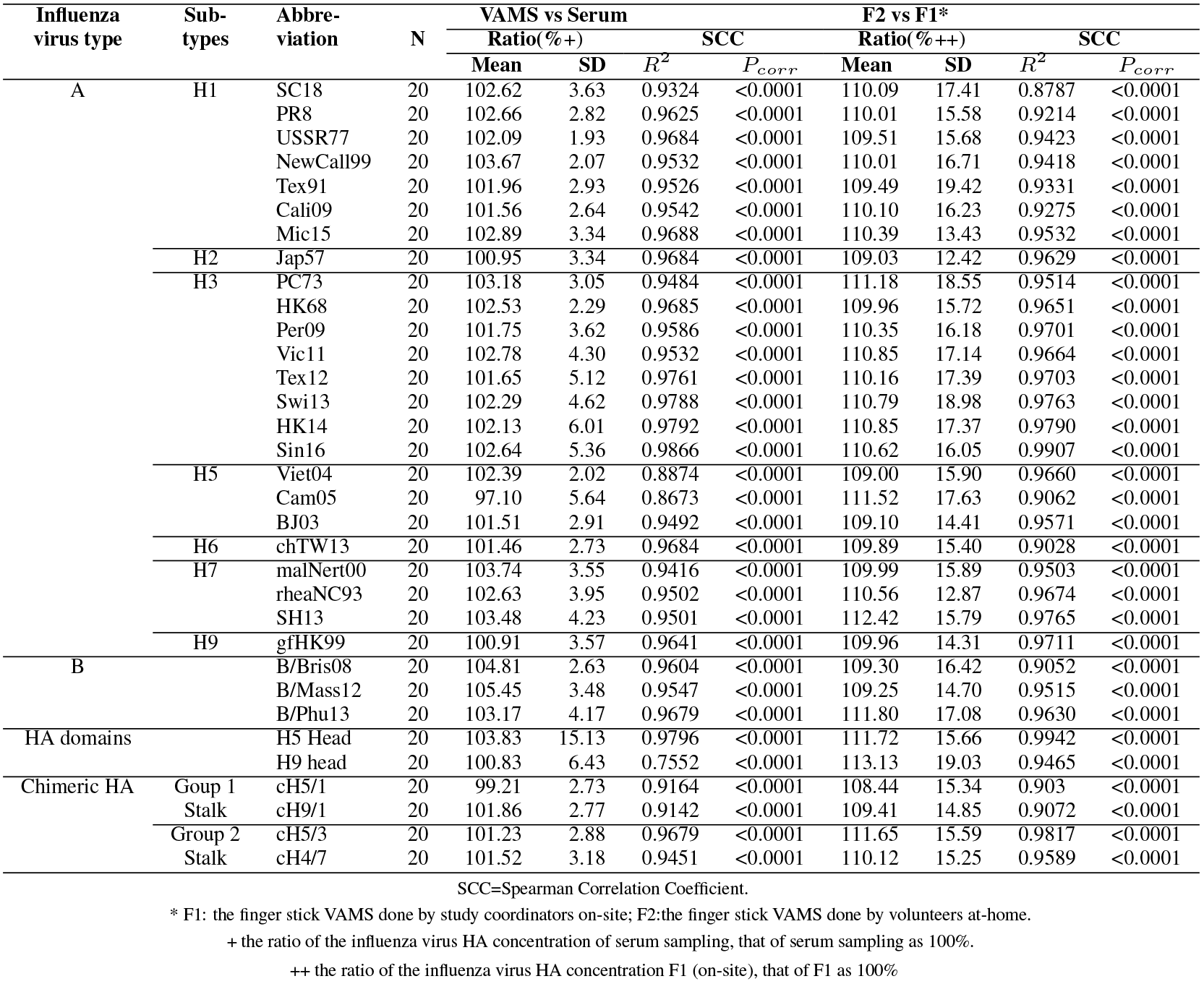
Correlation between mPLEX-Flu anti-HA IgG results: paired samples comparing VAMS vs. serum samples, and on-site versus remote VMAS sampling.

Using the same methods, we tested the correlation of mPlex-Flu results from VAMS and venupuncture sampling for 33 influenza virus strains, over the total 660 concentration data. We found a high correlation (*R*^2^ = 0.972; P <0.001) between the two methods (Figure 3). We found a similar correlation between serum sampling and VAMS when the correlation coefficients were calculated using the mean concentrations of anti-HA antibody, separated by subject, than when separated by influenza virus strains (mean *R*^2^ = 0.947), shown in Table 3. These results suggest that there is also a high correlation of VAMS with serum sampling for the mPlex-Flu assay to assess the individual influenza virus strains in each human subject. We found that the within-subject correlations were slightly higher than correlations across subjects (*R*^2^ range = 0.9310-0.9977), shown in Figure 4, due to smaller within-subject variability than between-subject variability. Both high within-subject and between-subject correlations provides strong evidence to support the high reliability of the VAMS method.

**Figure 3.**
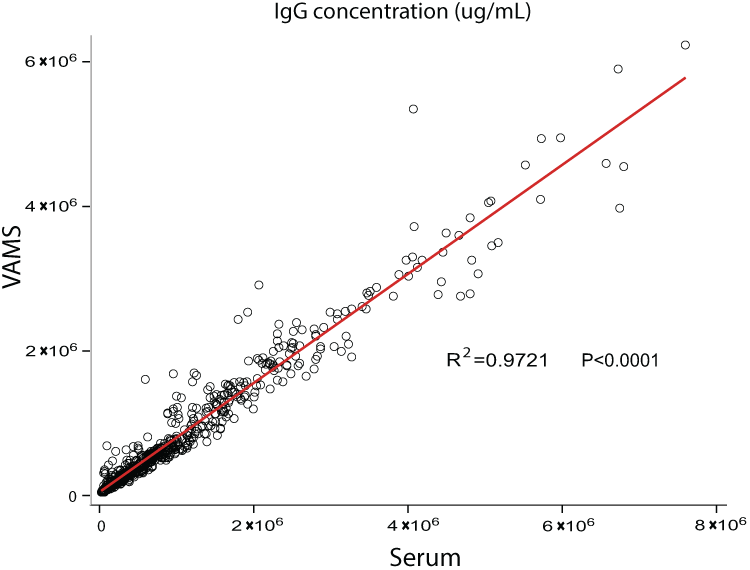
The overall correlation of concentration of influenza virus IgG antibodies against 33 strains of influenza virus by mPlex-Flu assay using VAMS sampling verse venous serum sampling. (n=660).

**Figure 4.**
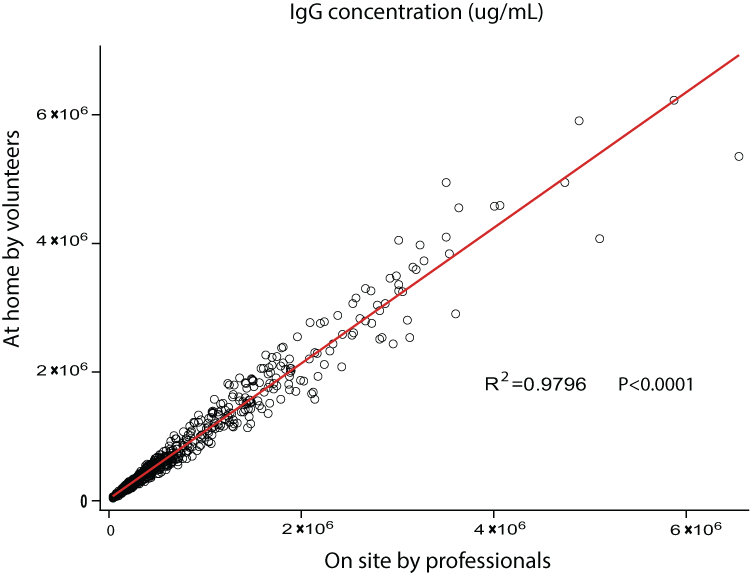
The overall correlation of concentration of influenza virus IgG antibodies against 33 strains of influenza virus by mPlex-Flu assay with VAMS finger stick from on site professionals with that from volunteers at home. (n=660)

### 3.3 VAMS is a highly reproducible process and can be performed at home

One advantage of the VAMS method is the safety and simplicity of the process. It is easy for study volunteers to learn and perform at home. Previously published data have shown that the volume of blood captured in the 10*µL* Matrix device varies <0.4*µL* [20]. But no study has shown the reproducibility of VAMS sampling by participants at home compared to on site by a nurse in the mPlex-Flu assay to evaluate influenza virus antibodies. To estimate this correlation, the same 20 participants also performed a second finger stick collection on themselves three days later. These samples were then hand-delivered back to the laboratory in a provided envelope.

Using the same analytic approach, we calculated the correlation of at-home (*F*2) and on-site (*F*1) sampling for measurement of IgG mediated immunity across multiple influenza virus strains, grouped by strain and subject. The results are shown in Table 3, and Figures 4 and 5. We found no statistically significant difference between the results obtained with on-site versus at-home VAMS sampling. This data suggests that VAMS sampling could be preformed at home by the study subjects, as the anti-HA antibody concentrations are highly consistent with sampling performed by study personnel on-site. These results support the consistency of VAMS sampling for future influenza vaccine or infection immunity studies.

**Figure 5.**
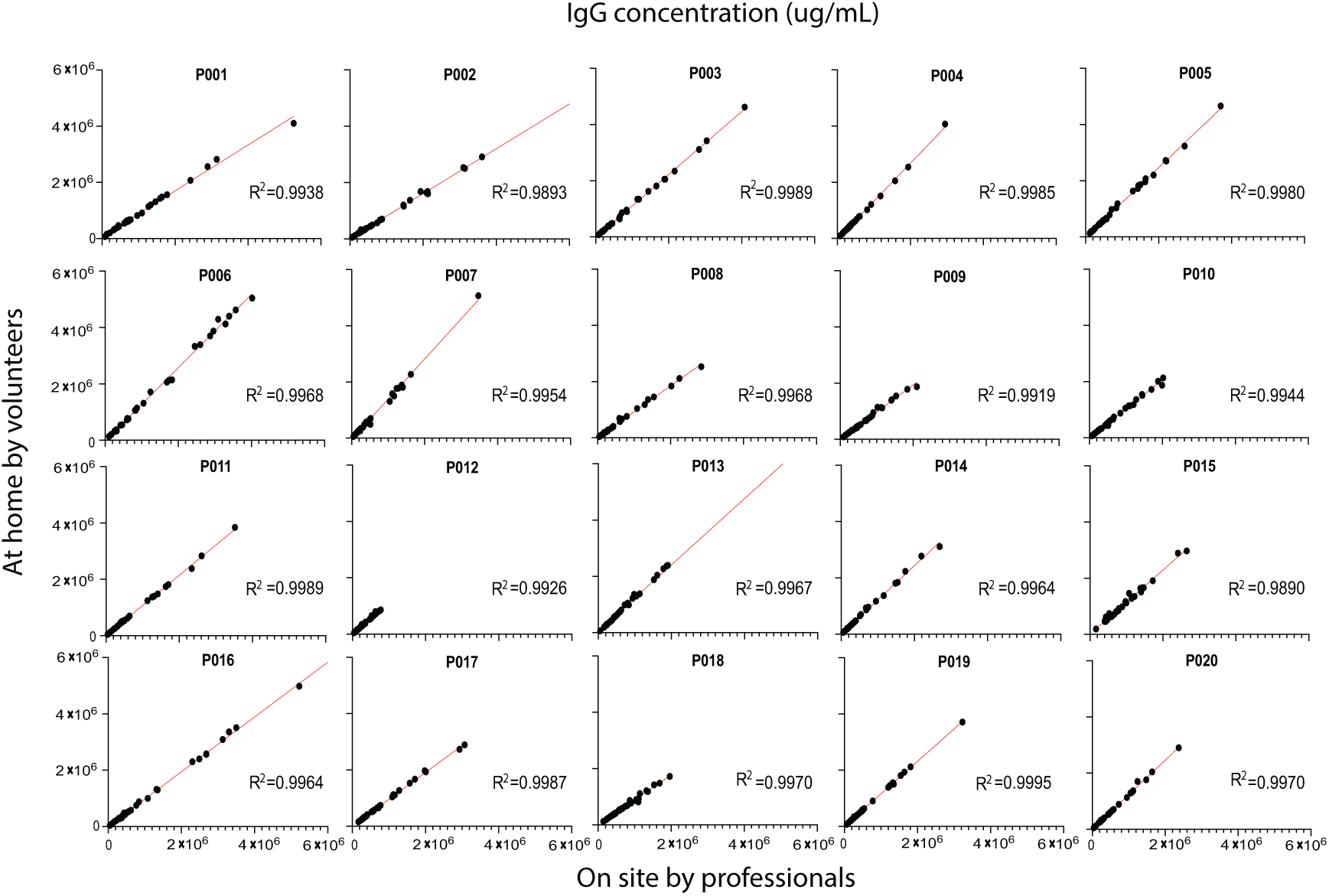
The correlation of concentration of influenza virus IgG antibodies against 33 strains of influenza virus by mPlex-Flu assay collected with VAMS finger stick from on site professionals with that from volunteers at home separated by individual subject (n=33).

### 3.4 Long-term stability of influenza virus HA antibodies in VAMS sampling device

We next examined the stability of anti-influenza virus HA antibodies in samples stored in VAMS device at room temperature, and during transport (e.g. postal service, two-day express mail). This is an essential aspect of quality control that needs to be addressed for future applications of VAMS. Prior studies have shown that antibodies on DBS filter paper are stable for more than 20 years when stored at 4°C or −20°C[19]. To determine the stability of antibodies in the VAMS device over time at room temperature, we used the mPlex-Flu assay to compare the antibody activity of VAMS tips stored at −20°C immediately after drying (control) with other VAMS tips left for 7, 14, 21 or 28 days at room temperature. The results are shown in Figure 6. We found no detectable antibody activity decrease at room temperature from storing the VAMS devices at room temperature environment, and antibody activities were still kept at 94.5% of the control level up to 21 days. After storing VAMS devices at room temperature to 28 days, the antibody activity level was significantly decreased to 80% of the control levels from the control devices stored at −20°C (statistical results were shown in table4).

**Figure 6.**
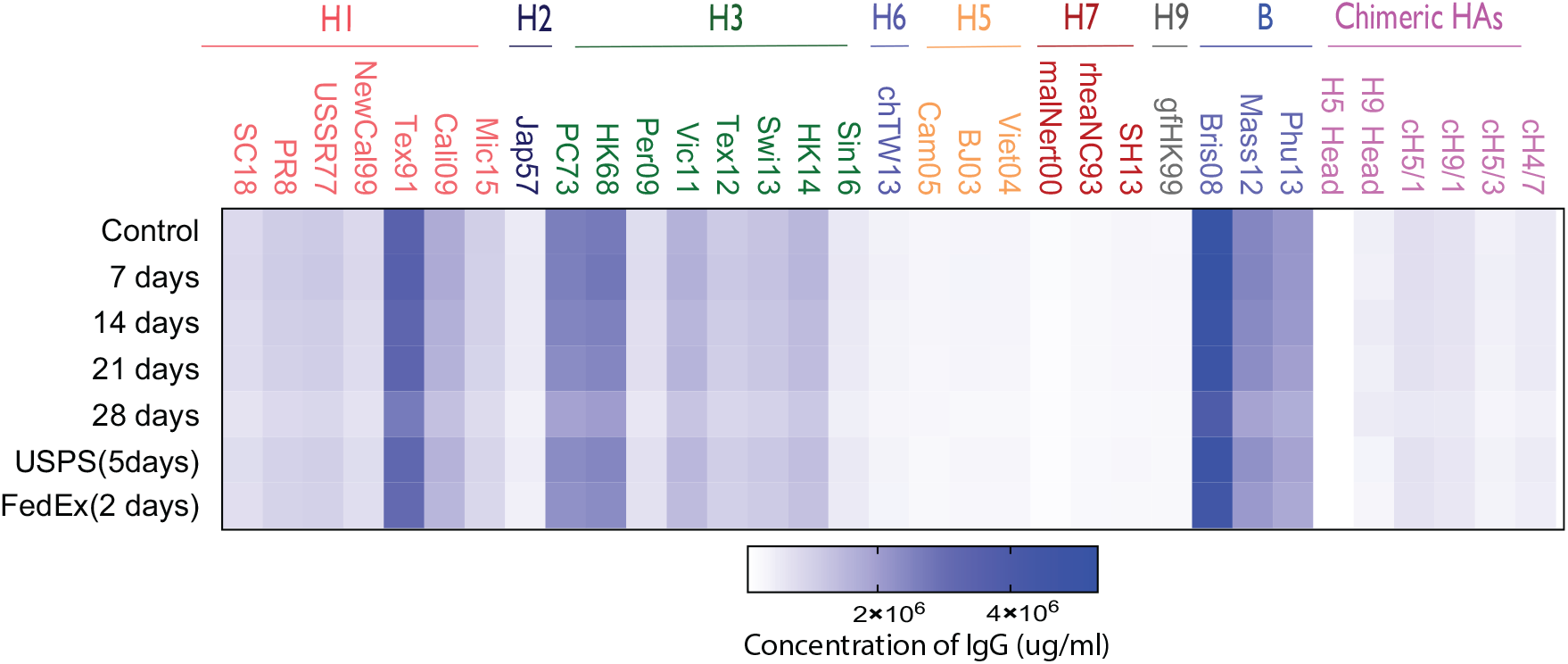
The stability of multiple dimensional IgG antibody with VAMS finger stick stored at room temperature or after shipping. The mean concentration of influenza virus IgG antibodies against 33 strains of influenza virus by mPlex-Flu assay shown in the heatmap. (n=4)

To confirm the stability of the antibodies during the shipping process, we compared two commonly used shipping methods: The United States Postal Service (USPS) First Class mail, and two-day commercial shipping (Federal Express, FedEx) to send two duplicate groups of the VAMS devices to our lab in New York State in August. USPS first class mail took 5 days, and commercial shipping took 2 days. After the samples were received back at the lab, the anti-HA IgG antibody levels were evaluated by mPlex-Flu assay (Figure 6). No statistically significant difference was detected between results from samples transported via the two shipping methods (Table 4), suggesting that the VAMS samples are stable during shipping process (2-5 days) even during the summer time, when temperatures may be elevated.

**Table 4.** The statistical results of VAMS specimen stability testing

| | Ratio of Control% | | Comparison to Control | | Correlation Adjusted $R^2$ |
| --- | --- | --- | --- | --- | --- |
|  | Mean | SD | t value | P value |  |
| Control | 100.00 | - | - | - | - |
| 7 days | 101.63 | 2.65 | -0.097 | 0.923 | 0.9828 |
| 14 days | 97.21 | 3.94 | -0.611 | 0.542 | 0.9306 |
| 21 days | 94.50 | 4.13 | -0.764 | 0.445 | 0.9835 |
| 28 days | 80.08 | 8.52 | **-2.267** | **0.024** | 0.9079 |
| Postal Service (5 days) | 91.23 | 3.22 | -1.087 | 0.278 | 0.9496 |
| Express (2 days) | 87.11 | 3.85 | -1.603 | 0.110 | 0.8936 |

Significant results from the generalized linear model in **bold text**.

## 4 Conclusions

We have demonstrated the utility of capillary VAMS sampling, combined with the mPlex-Flu assay, for measuring anti-influenza HA IgG antibody levels. The VAMS sampling method is inexpensive, can be used remotely by study volunteers, and yields consistent results compared to standard phlebotomy sampling. Finally, we found that VAMS samples are stable at room temperature for up to 21 days, and during the a standard 2-5 day shipping process.

The combination of VAMS sampling and mPlex-Flu analysis has the potential to greatly improve population studies of antibody mediated influenza immunity, vaccine response monitoring, and to augment the data collected by influenza surveillance field teams. For example, a major issue in vaccine trials is the expense and difficulty of having study subjects come to a study center for phlebotomy to monitor vaccine responses; a combined VAMS + mPlex-Flu methodology could allow for longer term remote sample collection from study subjects. Similarly, providing WHO or CDC teams with VAMS collection kits when surveilling influenza cases would permit collection of samples that could be analyzed to determine if infected subjects had antibody mediated immunity against circulating influenza strains. Furthermore, combining the antigenic distance of HA between different influenza strains via sequence comparisons, with the anti-HA IgG levels obtained by mPlex-Flu study of very large populations, could be used to create large-scale HA antigenic landscapes[24, 25] for future influenza virus infection and vaccine studies.

## 6 Acknowledgment

We would like to thank all the study volunteers, without whom this work would not be possible. We also are very grateful for the outstanding study coordination by Susanne Heininger, study coordinator, and the excellent technical assistance from the research coordinators at the University of Rochester Clinical Research Center.

This work was supported by the National Institutes of Health Institute of Allergy, Immunology and Infectious Diseases grant R21 AI138500 (MZ, JW), and the University of Rochester Clinical and Translational Science Award UL1 TR002001 from the National Center for Advancing Translational Sciences of the National Institutes of Health (JW, DL, MZ). The content is solely the responsibility of the authors and does not necessarily represent the official views of the National Institutes of Health. None of the above funders had any role in study design, data collection and analysis, decision to publish, or preparation of the manuscript.

## 7 Highlights

- First report to apply volumetric microsampling(VAMS) technique to influenza virus specific antibody assay
- Antibody levels of anti-influenza HA collected by VAMS highly correlate with conventional serum sampling in mPlex-Flu assay
- VAMS with mPLEX-Flu multiplex assay is a highly reproducible sample collection and analytic approach
- Influenza virus specific antibody blood samples are stable at room temperature and during shipping, and samples can be stored long term at 4°C
- Combining VAMS sampling and the mPlex-Flu assay provides a powerful tool for influenza virus antibody response studies

